# Multilayered interaction between CRISPR-Cas subtype I-A and adjacently encoded Acrs of virus SIRV2

**DOI:** 10.1101/2024.04.02.587611

**Authors:** Yuvaraj Bhoobalan-Chitty, Nicodemus Dwiputra, David Mayo-Muñoz, Karen Baadsgaard, Mette Rehtse Kvistrup Skafte Detlefsen, Xu Peng

## Abstract

Among the >100 anti-CRISPRs (Acrs) identified to date, the vast majority inhibit CRISPR-Cas immunity on its own. Here we report a multilayered interaction between CRISPR-Cas subtype I-A immunity and two Acrs encoded adjacently in the genome of Saccharolobus virus SIRV2, gp47 (AcrIA3) and gp48 (previously known as AcrIIIB1, hence termed AcrIIIB1/AcrIA4). The host subtype I-A CRISPR-Cas interference module was shown previously to be up-regulated upon SIRV2 infection, through the release of transcriptional repressor Csa3b from the promoter. We demonstrate that AcrIIIB1/AcrIA4 on its own increases viral infectivity 4-5 orders of magnitude in the presence of the host subtype I-A CRISPR-Cas immunity. This Acr is able to completely inhibit the subtype I-A CRISPR-Cas immunity when the transcriptional activation of the latter is artificially removed, suggesting that Acrs might be one of the driving forces for the evolution of CRISPR-Cas up-regulation. Interestingly, AcrIA3 cooperates with AcrIIIB1/AcrIA4 by inhibiting transcriptional activation of the host subtype I-A CRISPR-Cas interference module through interaction with the promoter of the latter. Taken together, our data shed light on how virus-host arms race shaped the evolution of CRISPR-Cas and Acrs.

## INTRODUCTION

CRISPR-Cas (Clustered Regularly Interspaced Short Palindromic Repeats – CRISPR associated genes) serve as the only known adaptive immune system that targets invading MGEs (Mobile Genetic Elements) in bacteria and archaea. Among the diverse CRISPR-Cas systems, the class 1 CRISPR-Cas, divided into type I, type III and type IV, have a high degree of complexity in terms of their RNA-guided surveillance complex. Subtype I-A CRISPR-Cas systems are widespread in archaea and consist of an interference and spacer acquisition module, both of which have been shown to be active against plasmids and virus infections (1-5). Notably, it has been shown in *Sulfolobales* that in response to viral infection the CARF (CRISPR Associated Rossman Fold)/HTH (Helix turn helix)-domain containing regulatory proteins Csa3a (activator of spacer acquisition module) and Csa3b (repressor of the interference module) are transcriptionally regulated to enhance CRISPR-Cas activity (6, 7).

The more than 100 known anti-CRISPRs (Acrs) exhibit diverse inhibitory mechanisms, including crRNA cleavage (8, 9), DNA mimicry and blocking of target DNA recognition (10-14), restriction of target entry (15), interference with complex biogenesis (16) or complex turnover (17), ADP-ribosylation (18) or acetylation (19) of Cas proteins, CRISPR repeat mimicking (20) and redirection of CRISPR-Cas complex to non-specific DNA (21) among others. Recently a AmrZ homolog, capable of transcriptional repression of CRISPR-Cas subtype I-F was identified on *Pseudomonas aeruginosa* mobile genetic element (22). The host encoded AmrZ is involved in regulation of the *P. aeruginosa* alginate biosynthesis pathway and was also shown to repress CRISPR-Cas subtype I-F expression (23). Despite inhibitors of most CRISPR-Cas systems having been identified, an Acr targeting the prevalent and well-studied subtype I-A interference was only recently identified, whose inhibitory mechanism remains however to be identified (24) .

The genome of *Sulfolobus islandicus* LAL14/1 encodes a subtype I-A, a subtype I-D and two subtype III-B CRISPR-Cas systems alongside two CRISPR arrays with subtype I-A specific repeats and three CRISPR arrays with subtype I-D specific repeats. Although inhibitors of subtypes I-D and III-B have been identified from the genome of Sulfolobus islandicus rod-shaped virus 2 (SIRV2), (25, 26), a lytic virus capable of infecting *S. islandicus* LAL14/1, an inhibitor of subtype I-A is yet to be identified from SIRV2.

In this study, we identified two adjacently encoded SIRV2 genes, gp47 and gp48 (previously identified AcrIIIB1 (26)) that collaboratively inhibit the subtype I-A system. Moreover, we show that gp47 (AcrIA3) transcriptionaly represses subtype I-A Cas gene expression. It carries a winged helix-turn-helix (wHTH) DNA binding motif that is structurally akin to the host subtype I-A transcriptional repressor Csa3b, AcrIA3 likely counters viral infection induced de-repression (i.e., activation) of the Cas operon maintaining it at a subdued pre-infection level.

## RESULTS

### SIRV2 gene product gp47 and gp48 inhibit CRISPR subtype I-A

A SIRV2 mutant, SIRV2M_II_, lacking the genes *gp02*-*gp09, gp51*-*gp53, gp10*-*gp14* and *gp48*, was susceptible to CRISPR-Cas targeting in *S. islandicus* LAL14/1 Δ*arrays*ΔIII-BΔI-D (a host carrying only subtype I-A CRISPR-Cas system) with a plasmid-borne mini-array containing the spacer *Sp*_*gp39*_, referred to as I-A strain (Figure 1A). In comparison to SIRV2M (lacking *gp02*-*gp09, gp51*-*gp53*), a decrease in infectivity of 4-5 orders of magnitude was observed with SIRV2M_II_. As expected, SIRV2M_II_ lost completely the infectivity in a host encoding subtype III-B only (Figure 1A**) (26)**. Loss of infectivity observed in case of the I-A strain is most likely due to the absence of *gp48* i.e., *acrIIIB1*.

**Figure 1.**
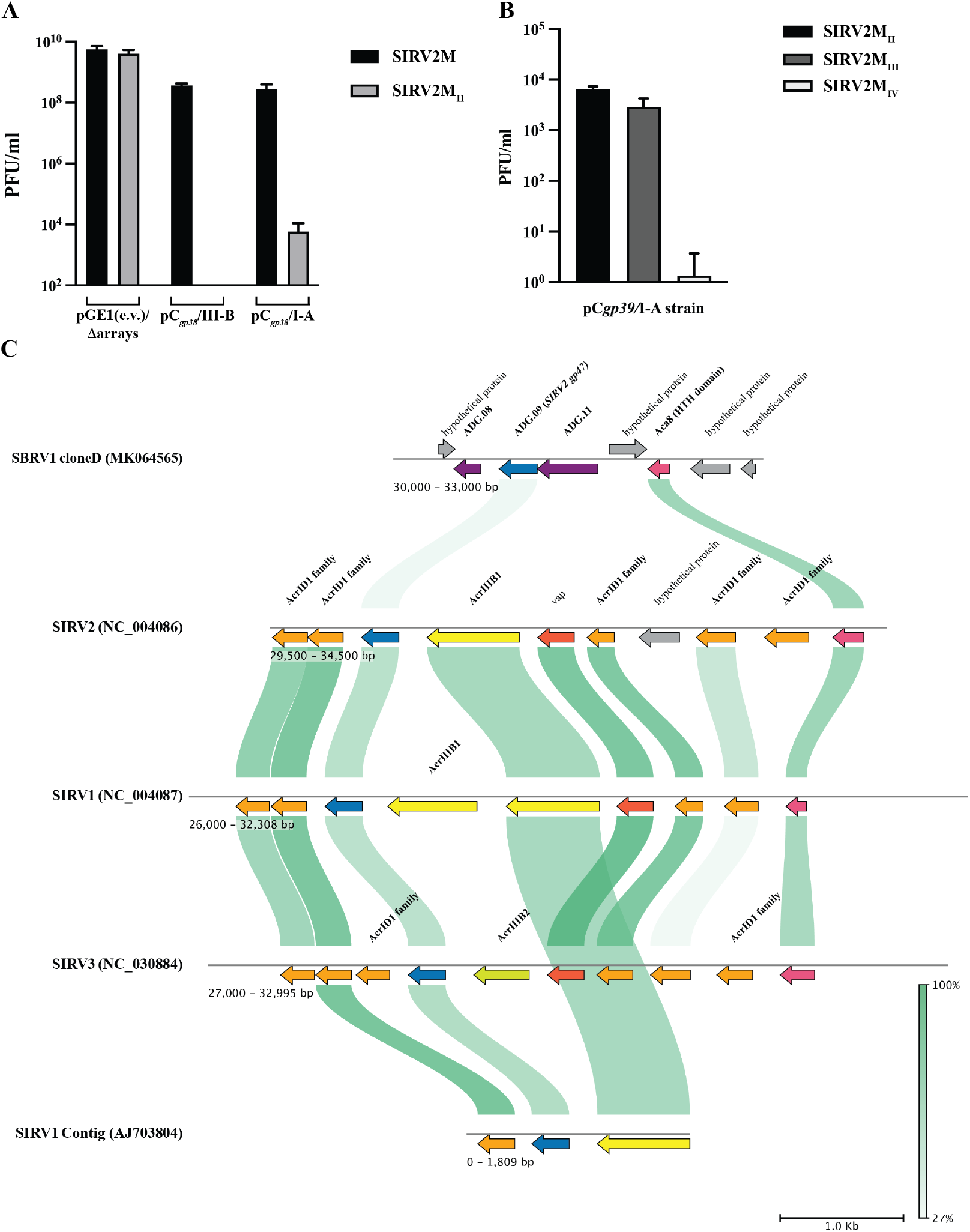
Inhibitors of subtype I-A in SIRV2. **A**. Plaque-forming efficiency of SIRV2M and SIRV2MΔgp48 in *S. islandicus* LAL14/1 Δ*arrays* carrying a spacer free vector (pGE1 (e.v. – empty vector)) in comparison with *S. islandicus* LAL14/1 Δ*arrays* Δ*I-A* Δ*I-D* (III-B strain) and *S. islandicus* LAL14/1 Δ*arrays*Δ*III-B*Δ*I-D* (I-A strain) both carrying the spacer Sp_*gp39*_. Data shown are mean of three biological replicates, represented as mean ± SD. **B**. Plaque-assay of SIRV2MII mutants in *S. islandicus* LAL14/1 Δ*arrays* I-A strain with the plasmid-borne spacer Sp_*gp39*_. **C**. Genomic location of SIRV2 gp47 homologs among different archaeal viruses. AcrIIIB1/AcrIA4 homologs (in blue) and its neighboring anti-defense genes (ADGs) in genomes of archaeal viruses. AcrIIIB1 homologs in yellow, AcrID1-family proteins in orange, Aca8 in dark pink, hypothetical proteins in grey and other predicted SBRV1 ADGs, ADG.08 and ADG.11 in purple.

A recent bioinformatic study predicted several hundred anti-defense genes (ADGs) based on regulatory sequences in archaeal viral genomes (24). Apart from previously known Acrs such as AcrIIIB1, AcrID1 and AcrID1 homologs, SIRV2 gp47 (ADG.09) was among others predicted to be a putative ADG. Previously, SIRV2M_II_ was subjected to successive deletions of accessory genes leading to the construction of five additional mutants, SIRV2M_III_ – SIRV2M_VII_ (27). Among these, SIRV2M_IV_, but not SIRV2M_III_ was unable to propagate at all in the I-A strain (Figure 1B). Given the complete loss of infectivity upon deletion of *gp45*-*gp47* and the close association of *gp47* with *acrIIIB1* (*gp48)* and *acrID1* homologs (*gp45* and *gp46*), we inferred that gp47 is a likely AcrIA (Figure 1C).

To verify the role of SIRV2 gp47 through plasmid complementation assays, we transformed spacer containing plasmids encoding either *gp47* or the co-transcribed *gp46*-*gp45* under the control of an arabinose promoter into strains with or without subtype I-A interference *cas* genes. The transformation efficiency of *gp46*-*gp45* plasmid was nearly identical in both host strains but a ∼100-fold decrease in transformation efficiency was observed with the *gp47* plasmid in the I-A strain in comparison to the strain lacking subtype I-A (Figure 2A). The toxicity observed exclusively in the presence of I-A CRISPR-Cas was a strong indication of interaction between gp47 and I-A CRISPR-Cas in *S. islandicus* LAL14/1. Before conducting the complementation assay, we deleted *gp45*-*gp47* from the SIRV2 mutants SIRV2M and SIRV2M_II_ because the latter virus lacked *gp48* in comparison to the former. The two parental and two deletion mutants are hereafter referred to as SIRV2M, Δ*gp48*, Δ*gp45-gp47* and Δ*gp45*-*gp48* (Supplementary figure 1). Complementation assays were performed in I-A strains carrying plasmid-borne *gp48* or *gp47* with spacers *Sp*_*gp38*_ and *Sp*_*gp39*_, respectively, whose corresponding protospacers are located on all virus mutants utilized in this study. Following infection, complete growth retardation, indicative of CRISPR-Cas inhibition and virus propagation, was observed only in the presence of both *gp47* and *gp48*, either plasmid-borne or virus encoded (Figure 2B). Notably, mild growth retardation was observed upon infection of I-A strain carrying spacer *Sp*_*gp38*_ with virus Δ*gp48*, i.e., only gp47 was present (Figure 2B, left panel) but no growth retardation was observed upon infection of I-A strain carrying spacer *Sp*_*gp39*_ with virus Δ*gp45-gp47*, i.e., only gp48 was present (Figure 2B, right panel), suggestive of a stronger inhibition by gp47 alone than by gp48 alone. Taken together, we conclude that both SIRV2 gp47 and SIRV2 gp48 are inhibitors of subtype I-A and henceforth will be referred to as AcrIA3 and AcrIIIB1/AcrIA4, respectively. Furthermore, we can also infer that complete inhibition of *S*. islandicus LAL14/1 subtype I-A CRISPR-Cas system requires cooperation between both the Acrs.

**Figure 2.**
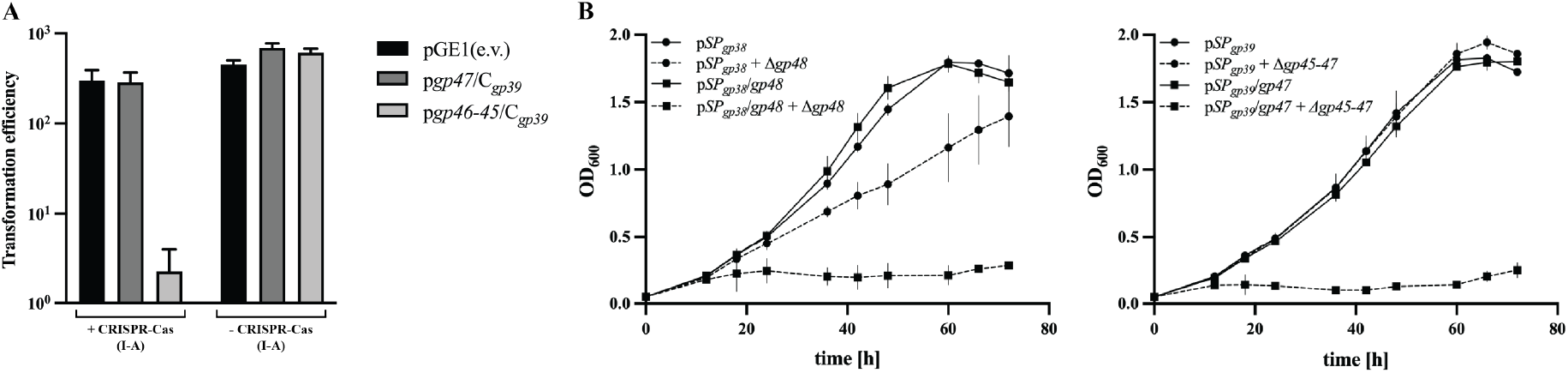
SIRV2 gp47 and SIRV2 gp48 co-operatively inhibit CRISPR-Cas subtype I-A in *S. islandicus* LAL14/1. **A**. Transformation efficiency of plasmids carrying *gp46*-gp*45* (red) or *gene47* (green) alongside Sp_*gp39*_ spacer carrying mini-CRISPR array in *S. islandicus* LAL14/1 Δ*arrays*/ΔIII-B strain with or without subtype I-A system. Data shown are mean of three biological replicates, represented as mean ± SD. **B**. Complementation assay in Δ*arrays*/ΔIII-B strain carrying the spacer encoding plasmids pSp_*gp38*_(left panel)/pSp_*gp39*_(right panel) without (filled circle) or with (filled square) *gene48* (left panel) / *gene47* (right panel). Cells were infected with SIRV2MΔ*gp48* (left panel) or SIRV2MΔ*gp45*-*47* (right panel), infected culture are shown in dashed lines.

### AcrIA3 diminishes CASCADE transcripts

Previous transcriptomic study of *S. islandicus* LAL14/1 observed a 10-fold increase in the expression of CASCADE operon (subtype I-A Cas genes) upon infection with SIRV2 (28). A subsequent study showed that P_*cas*_ (regulatory and promoter sequence of the CASCADE operon) is repressed cooperatively by the subtype I-A CASCADE complex and Csa3b. Moreover, P_*cas*_ is derepressed by about 40- and 25-fold in the absence of CASCADE and Csa3b, respectively (7). The discrepancy between the potential maximum derepression (40-fold) and the lower derepression during virus infection (10-fold) led us to consider the possibility that one or both identified Acrs could be involved in transcriptional regulation.

The transcript levels of *csa5* (first gene in the CASCADE operon) were estimated in *S. islandicus* Δ*arrays* strains carrying empty vector, plasmid-borne AcrIIIB1/AcrIA4 or AcrIA3, either with or without virus infection by Δ*gp45*-*gp48*. Whereas *acrIIIB1*/*acrIA4* showed no big effect on *csa5* transcript level, the plasmid-borne *acrIA3* strongly inhibited transcription of *csa5*, as shown by the >10,000 fold less *csa5* in comparison to that measure in cultures containing the empty plasmid (Figure 3A and Supplementary figure 2C). These results confirm that there is a decrease in the CASCADE transcript count in the presence of AcrIA3.

**Figure 3.**
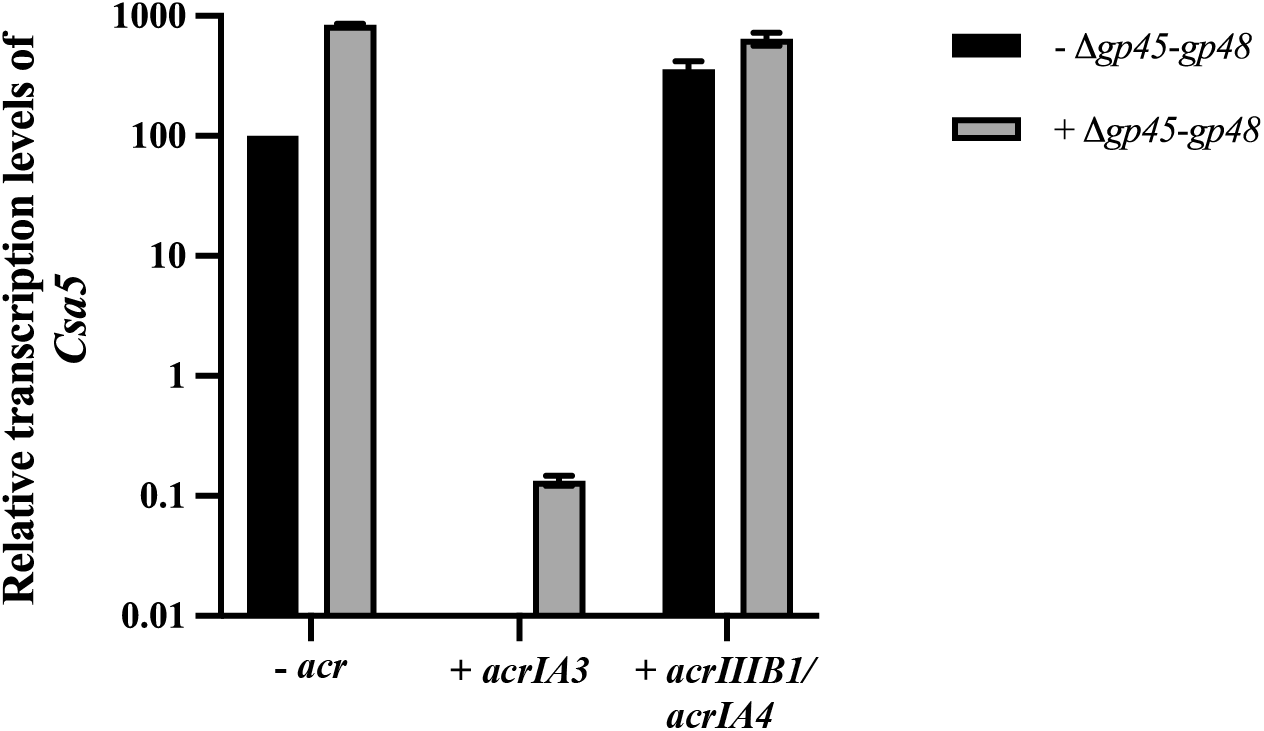
AcrIA3 downregulates CASACDE transcription. **A**. qRT-PCR measurement of relative transcription levels of *csa5*, representing CASCADE transcription, in cultures without Acr (-*acr*) or in the presence of either AcrIA3 (+ *acrIA3*) or AcrIIIB1/AcrIA4 (+ *acrIIIB1/acrIA4*). Measurement of transcription levels was performed in the described cultures without (red) and with (green) virus infection. Data shown are average of values obtained from two independent qPCRs, represented as mean ± SD.

### AcrIA3 impedes derepression of CASCADE in a P_*cas*_ - dependent manner

Next, we aimed to understand the AcrIA3 dependent decrease in the *csa5* transcript levels observed earlier and to examine if it was due to transcriptional repression of P_*cas*_. To achieve this the Δ*arrays* strain was genetically modified to substitute P_*cas*_ with the synthetic arabinose inducible promoter, *ara*_*S2*_ (Supplementary figure 3), this strain was referred to as Δ*arrays*/P_*cas*_::P*ara*_*S2*_ (29). A plasmid carrying a mini-CRISPR array containing spacer *Sp*_*gp39-2*_, complementary to the template strand, was transformed into Δ*arrays*/P_*cas*_::P*ara*_*S2*_ and the wildtype Δ*arrays* strain carrying the native P_*cas*_ (hereafter referred to as as Δ*arrays*/P_*cas*_). The two strains were infected with SIRV2M and the Acr-deletion mutants, Δ*gp48*, Δ*gp45-gp47* and Δ*gp45*-*gp48*. Growth curves were plotted, and virus titers in culture supernatants were measured 12 and 24 h.p.i (hours post infection). Upon infecting the Δ*arrays*/P_*cas*_/p*Sp*_*gp39-2*_, only the parental virus SIRV2M carrying both AcrIAs was able to cause growth retardation (Figure 4A). Whereas upon infection of Δ*arrays*/P_*cas*_::P*ara*_*S2*_/ p*Sp*_*gp39-2*_, besides the parental virus, the virus Δ*gp45-gp47* was also able to cause growth retardation (Figure 4B), indicating that AcrIA3 might not be essential for infection in this host strain. Despite replacement of the promoter sequence, the inability of the viruses Δ*gp48* and Δ*gp45*-*gp48* to propagate in the Δ*arrays*/P_*cas*_::P*ara*_*S2*_/pC_*gp39*-2_ strain demonstrates subtype I-A’s effectiveness in countering virus propagation and that the inhibition mechanism of AcrIIIB1/AcrIA4 does not involve transcriptional repression of CASCADE. These findings were additionally validated through titration of virus in the supernatant, only SIRV2M was able to propagate efficiently in the wildtype host (Figure 4C) but both viruses encoding AcrIIIB1/AcrIA4 were able to propagate in the Δ*arrays*/P_*cas*_::P*ara*_*S2*_ host, regardless of the existence of AcrIA3 (Figure 4D).

**Figure 4.**
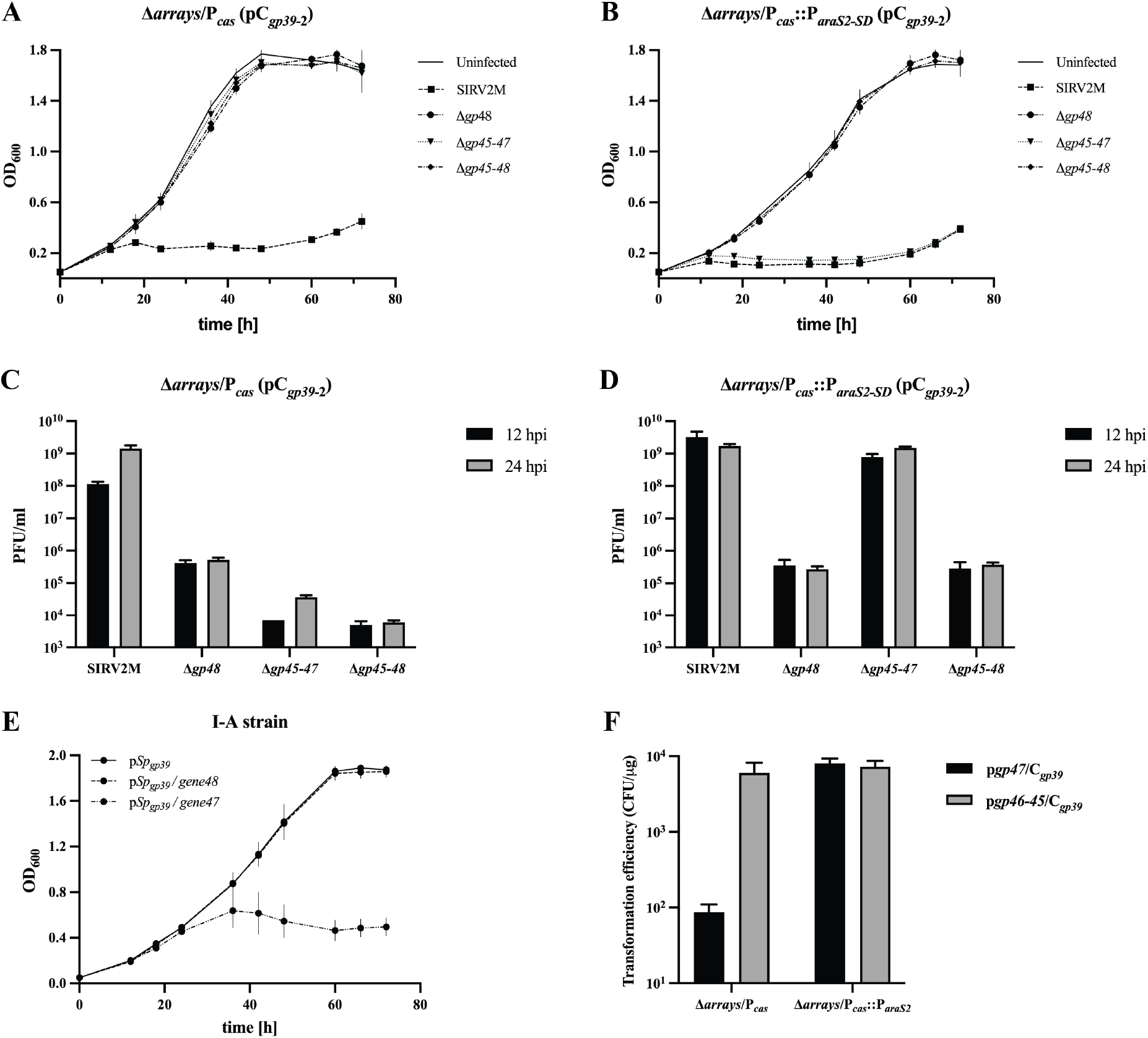
CASCADE repression by AcrIA3 is P_*cas*_ promoter dependent. **A**. and **B**. Growth curves of *S. islandicus* LAL14/1 Δ*arrays* culture carrying the native P_*cas*_ promoter and *S. islandicus* LAL14/1 Δ*arrays* culture carrying the synthetic P_*araS2*_ promoter respectively upon infection with SIRV2M, SIRV2MΔ*gp48*, SIRV2MΔ*gp45*-*47* and SIRV2MΔ*gp45*-*gp48*. The multiplicity of infection (MOI) = 3 (**A)** and 0.8 (**B**). Data shown are mean of three biological replicates, represented as mean ± SD. **C**. and **D**. Titration of virus quantity in cultures by plaque assay using samples from cultures defined in **A**. and **B**. obtained 12 and 24 hours post infection (h.p.i). Data shown are mean of three biological replicates, represented as mean ± SD. **E**. Growth curves of *S. islandicus* LAL14/1 Δ*arrays*/ΔIII-B strains carrying spacer (*Spgp39*) containing plasmids, without or with either *acrIA3* or *acrIIIB1*/*acrIA4*, upon infection with the Acr-free virus Δ*gp45-gp48*. **F**. Comparison of transformation efficiencies of plasmids carrying spacer C_*gp39*_ and *gp47* or *gp45*-*46*, in Δ*arrays*/P_*cas*_::P_*araS2*_ and Δ*arrays*/P_*cas*_.

To examine the importance of CRISPR-Cas downregulation to virus survival, I-A strains carrying spacer containing plasmid without or with either *acrIIIB1*/*acrIA4* or *acrIA3* were infected with the virus mutant lacking both Acrs, Δ*gp45*-*gp48*. Although virus infection did not cause significant growth inhibition as observed previously in the presence of both Acrs (Figure 2B, Figure 3A), the Acr-free virus caused delayed host growth retardation in the presence of the plasmid-borne AcrIA3 (Figure 4E), implying subtype I-A repression prior to infection sufficiently protects the virus from CRISPR-Cas targeting.

Earlier it was seen that *acrIA3* reduced transformation efficiency of the plasmid in the I-A strain, specifically in the presence of a functional subtype I-A effector complex (Figure 2A). To better understand this result we transformed plasmid-borne *acrIA3* along with the spacer *Sp*_*gp39*_ into Δ*arrays*/P_*cas*_ and Δ*arrays*/P_*cas*_::P*ara*_*S2*_ strains. Despite both strains carrying a functional subtype I-A CRISPR-Cas complex, plasmid-borne *acrIA3* was toxic in Δ*arrays*/P_*cas*_ (similar to the I-A strain) but not in Δ*arrays*/P_*cas*_::P*ara*_*S2*_ strain (Figure 4F). From these results it is clear that AcrIA3 downregulates CASCADE transcription by way of a mechanism which involves the subtype I-A promoter P_*cas*_.

### AcrIA3 is a HTH protein, similar to Sso Csa3b HTH domain

PSI-BLAST of AcrIA3 over three iterations (e-value = 0.05) retrieved only five viral homologs (Supplementary figure 5A and Supplementary figure 4B). To gain insights into the functionality of AcrIA3, we predicted its tertiary structure using AlphaFold2 (30) using default parameters. A winged helix-turn-helix structure was predicted for AcrIA3 in all five models with very high confidence (Figure 5A, Supplementary figure 5A and Supplementary figure 5B). Following this, we compared the predicted structure against known protein structures present in the Protein Data Bank (PDB) through the DALI server (31). Interestingly, structural similarity was observed with HTH regulatory proteins such as sarS (*Staphylococcus aureus*, 1P4X) (32), MarR family proteins (*Staphylococcus aureus*, 4XRF/5F6F) and Csa3b (*Sulfolobus solfataricus* P2, 2WTE) (33) (Figure 5B, Supplementary file 1). Based on this, we can infer that AcrIA3 is a virally encoded transcriptional regulator of P_*cas*_, akin to the subtype I-A associated regulator Csa3b.

**Figure 5.**
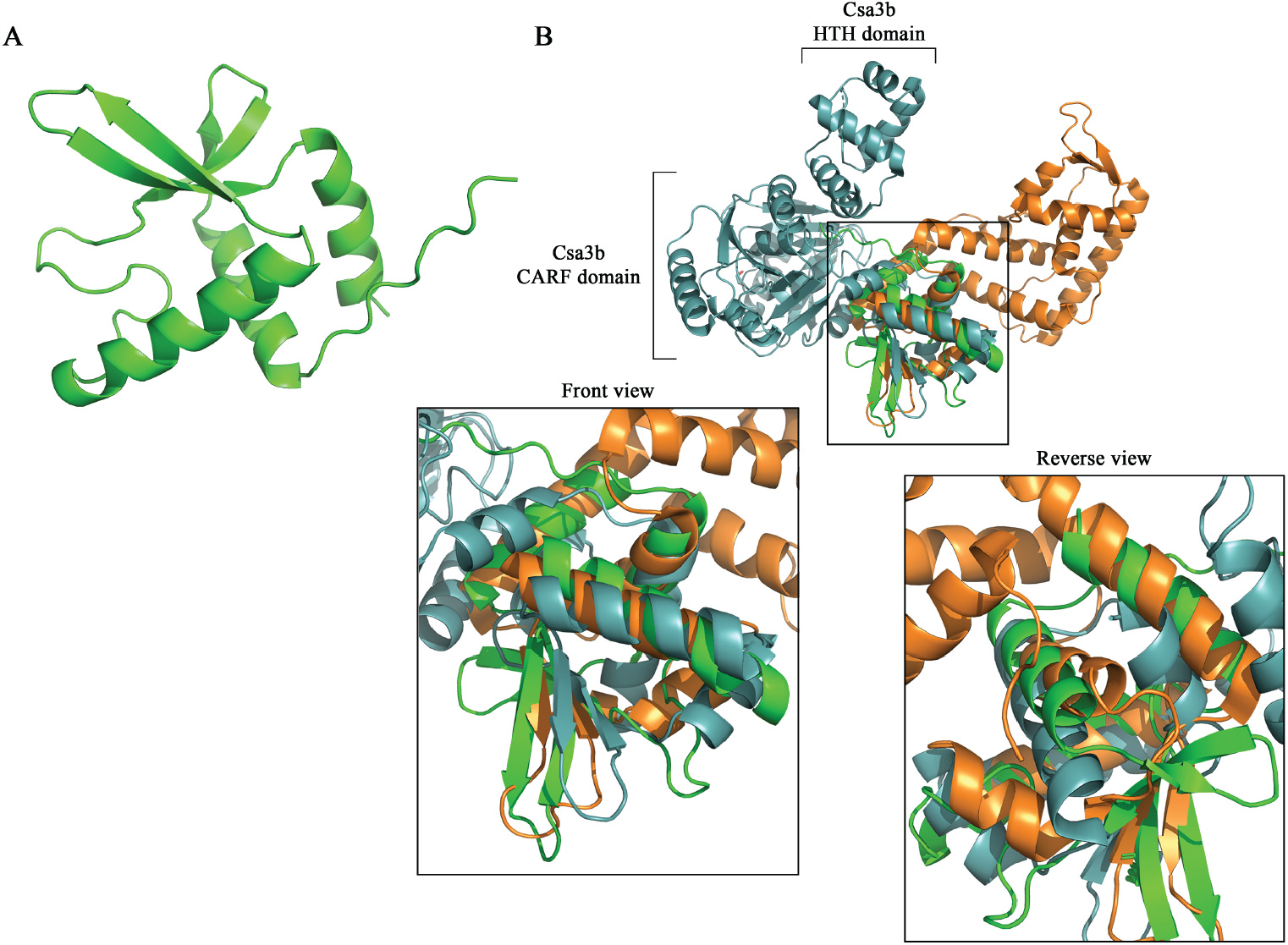
Prediction and comparison of AcrIA3 structure. **A**. AlphaFold2 structure prediction of SIRV2 gp47 (AcrIA3). **B**. Superimposition of predicted AcrIA3 structure (green) with SarS1 (1P4X, orange) and Csa3b homodimer (2WTE, deep teal), showing structural similarity between their HTH domains. Structural homologs were identified by comparison with protein structures in the Protein Data Bank using DALI server.

## DISCUSSION

Acrs, inhibiting the same CRISPR-Cas subtype, are only occasionally encoded within a single bacteriophage genome (8, 34). Depending on the inhibitory efficiency, Acr(s) dictate the initial viral density threshold required to establish complete host immunosuppression, Acrs with low efficiency requiring higher virus titer thresholds (35-37). Despite the absence of AcrIIIB1/AcrIA4 only resulting in a loss of infectivity by 4-5 orders of magnitude (Figure 1A), it is by itself not a weak Acr. As seen in a non-regulated state of the *Sulfolobus* subtype I-A, AcrIIIB1/AcrIA4 alone is sufficient to overcome CRISPR-Cas immunity (Figure 4B and Figure 4D). To counter typical Acrs (such as AcrIIIB1/AcrIA4) or possibly higher virus titres, *Sulfolobus* CRISPR-Cas subtype I-A derepress CASCADE expression by around 40-fold (7), depicting a likely anti-anti-CRISPR mechanism. The ability of Δ*gp45*-*gp47* to infect the Δ*arrays*/P_*cas*_::P*ara*_*S2*_ strain but not Δ*arrays*/P_*cas*_ strain demonstrates the potency of CASCADE derepression in overcoming virus infection despite the presence of AcrIIIB1/AcrIA4 (Figure 4A-4D). Consequently, this depression necessities viruses to infect at higher virus titers or evolve high-efficiency Acrs or instead evolve a novel inhibitory strategy, such as a repressor of CRISPR-Cas transcription as seen here akin to an anti-anti-anti-CRISPR, to overcome host immunity.

Similar to the *P. aeruginosa* AmrZ (22), AcrIA3 function as a transcriptional repressor of Cas gene expression, representing the most efficient strategy to overcome CRISPR-Cas derepression. Based on sequence similarity, AmrZ is likely to have host origins but purely structural similarity between SSOCsa3b and AcrIA3 means host origin is unlikely (Figure 5). Whereas AmrZ was shown to be active only during the lysogenic life cycle, AcrIA3 possess inhibitory activity during the lytic life cycle resulting in a mild host growth retardation (Figure 2B, left panel and Figure 4E). Three out of the five AcrIA3 homologs are encoded adjacent to AcrIIIB1/AcrIA4. SIRV3gp40 recently shown to be an inhibitor of subtype III-B (AcrIIIB2) is adjoining the AcrIA3 homolog in SIRV3 (SIRV3gp39) (17) (Figure 1C). Although it has not been tested for inhibitory activity against subtype I-A, it is likely to also posses subtype I-A inhibitory activity with efficiency similar to that of AcrIIIB1/AcrIA4. Furthermore, we expect that the AcrIA3 homolog (in SBRV1) and AmrZ to be effective only in co-operation with another inhibitor of subtype I-A (Figure 1C). Consequently, it is likely that AcrIIIB1/AcrIA4 homologs would also be flanked by other inhibitors (including transcriptional inhibitors) of subtype I-A as evidenced in SIFV and SIFV2 (24). Along with the *P. aeruginosa* phage encoded AmrZ homologs, AcrIA3 represents a novel family of regulatory Acrs which form vital elements in the host-virus evolutionary interplay.

## MATERIALS AND METHODS

### Strains and growth conditions

*S. islandicus* LAL14/1 Δ*arrays* and its derivatives were grown at 78°C, 150 rotations per minute in SCV medium, uracil (20μg/ml) was included when necessary. *Escherichia coli* DH5α was utilized for cloning of the *Sulfolobus*-*Escherichia* shuttle vector pEXA. All primers utilized in this study are listed in Supplementary table 1. *Sulfolobus* electroporation was performed as described earlier (38, 39).

### *Sulfolobus* and virus genetic manipulation

Replacement of P_*cas*_ promoter with P_*araS2*_ promoter was accomplished using the CRISPR-Cas dependent genome editing technique previously demonstrated in *Sulfolobus* (40). Briefly, 500bp homologous arms flanking the P_*cas*_ promoter were amplified and fused with the arabinose promoter. A spacer targeting the P_*cas*_ promoter was inserted into the mini-CRISPR array in pGE2 plasmid alongside the homologous arms with the desired insertion. The plasmid was electroporated into Δ*arrays* and transformants were screened for the correct insertion. 5-FOA (5-Fluororotic Acid, 50μg/ml) was used to cure the positive transformants of the genome editing plasmid. Deletion of SIRV2 *gp45*-*gp47* from SIRV2M and SIRV2M_II_ based on CRISPR-Cas genome editing method as described earlier (27).

### *Sulfolobus* growth assay and Virus plaque assay

Overnight *Sulfolobus* cultures were diluted to OD_600_ = 0.05 and infected at an MOI = 0.01, the cultures were sampled at specified timepoints for OD measurement and virus titration. Plaque assay for virus titration was done as described earlier (38).

### MSA and phylogenetic analysis

SIRV2 gp47 (NP_666581.1) homologs were identified by PSI-BLAST with a expect threshold of 0.05. In addition to data based on promoter sequence identification and comparison of homologs we deduced that translation initiation occurs 13 residues downstream of the predicted start codon. The homologs were aligned with Clustal Omega (41), a phylogenetic tree was constructed using neighbor-joining method and visualized using iTOL (42).

### SIRV2 gp47 structure and functional prediction

Truncated SIRV2 gp47 protein sequence, as described according to the positioning of the promoter sequence, was used as input for structure prediction in AlphaFold2 (default parameters) (https://colab.research.google.com/github/sokrypton/ColabFold/blob/main/AlphaFold2.ipynb).

The predicted model was used for comparison against structures in the Protein Data Bank using the DALI protein structure comparison server. All protein structures were visualized using PyMOL.

## DATA AVAILABILITY

Data supporting the findings in this study are available within the paper and its supplementary information. Additional data can be obtained from the corresponding author upon request.

## Author contribution statement

**Yuvaraj Bhoobalan-Chitty:** Conceptualization, Investigation, Formal analysis, Supervision, Validation, Funding acquisition, Writing – original draft, Writing – review and editing.

**Nicodemus Dwiputra**: Investigation, Formal analysis.

**David Mayo-Muñoz**: Investigation, Formal analysis.

**Karen Baadsgaard**: Investigation, Formal analysis.

**Mette Rehtse Kvistrup Skafte Detlefsen**: Investigation, Formal analysis.

**Xu Peng**: Conceptualization, Supervision, Validation, Funding acquisition, Writing – review and editing.

## ACKNOWLEDGEMENTS

All members of the Microbial Immunity Lab at the Copenhagen University are thanked for their suggestions. This work was supported by the Novo Nordisk Fonden Postdoctoral Fellowship in Bioscience and Basic Biomedicine Grant [NNF21OC0067491] to Y.B.-C., and by the Danish Council for Independent Research/Natural Sciences [DFF-0135-00402] and Novo Nordisk Foundation/Hallas Møller Ascending Investigator Grant [NNF17OC0031154] to X.P.

## SUPPLEMENTARY FIGURES

**Supplementary figure 1.**
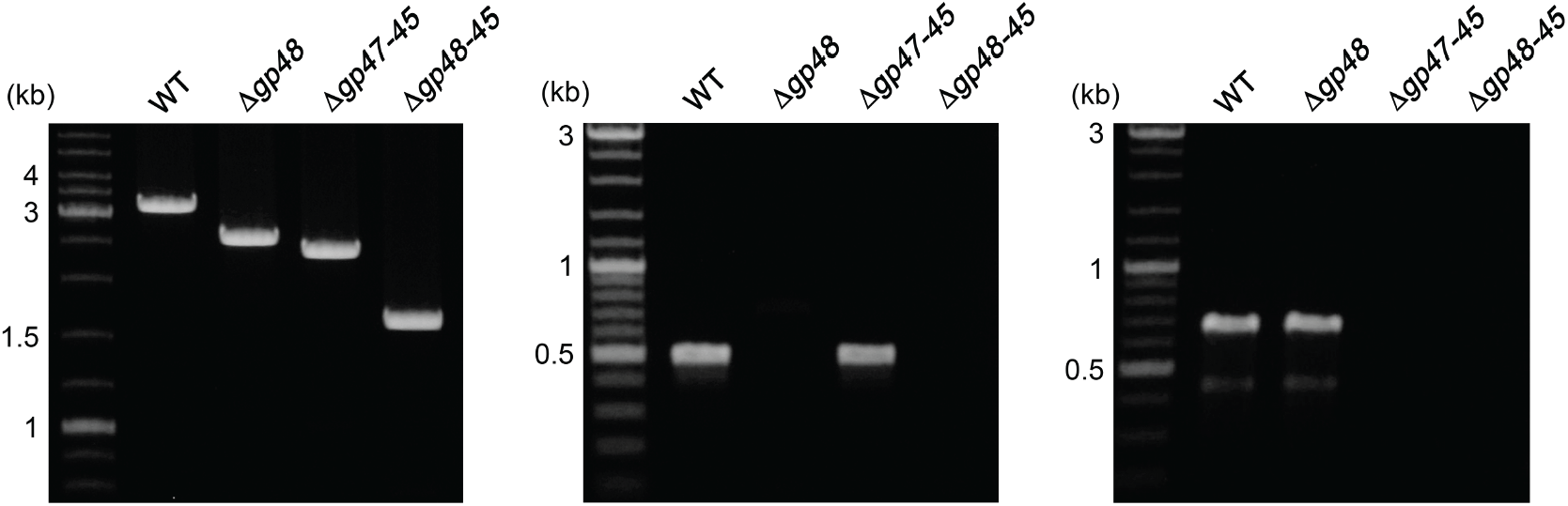
PCR verification of SIRV2M and its variants. The presence of the desired mutations in SIRV2M, SIRV2M_II_ (SIRV2MΔ*gp48*), SIRV2MΔ*gp45*-*gp47* and SIRV2M_II_Δ*gp45*-*gp47* (SIRV2MΔ*gp45*-*gp48*) was checked using PCR with virus supernatants as templates. Primers flanking the *gp45*-*gp48* (left panel), *gp48*-specific (middle panel) and *gp47*-specific (right panel) were used to check for the presence of the desired deletions. Primers utilized in this study are listed in Supplementary table 1.

**Supplementary figure 2.**
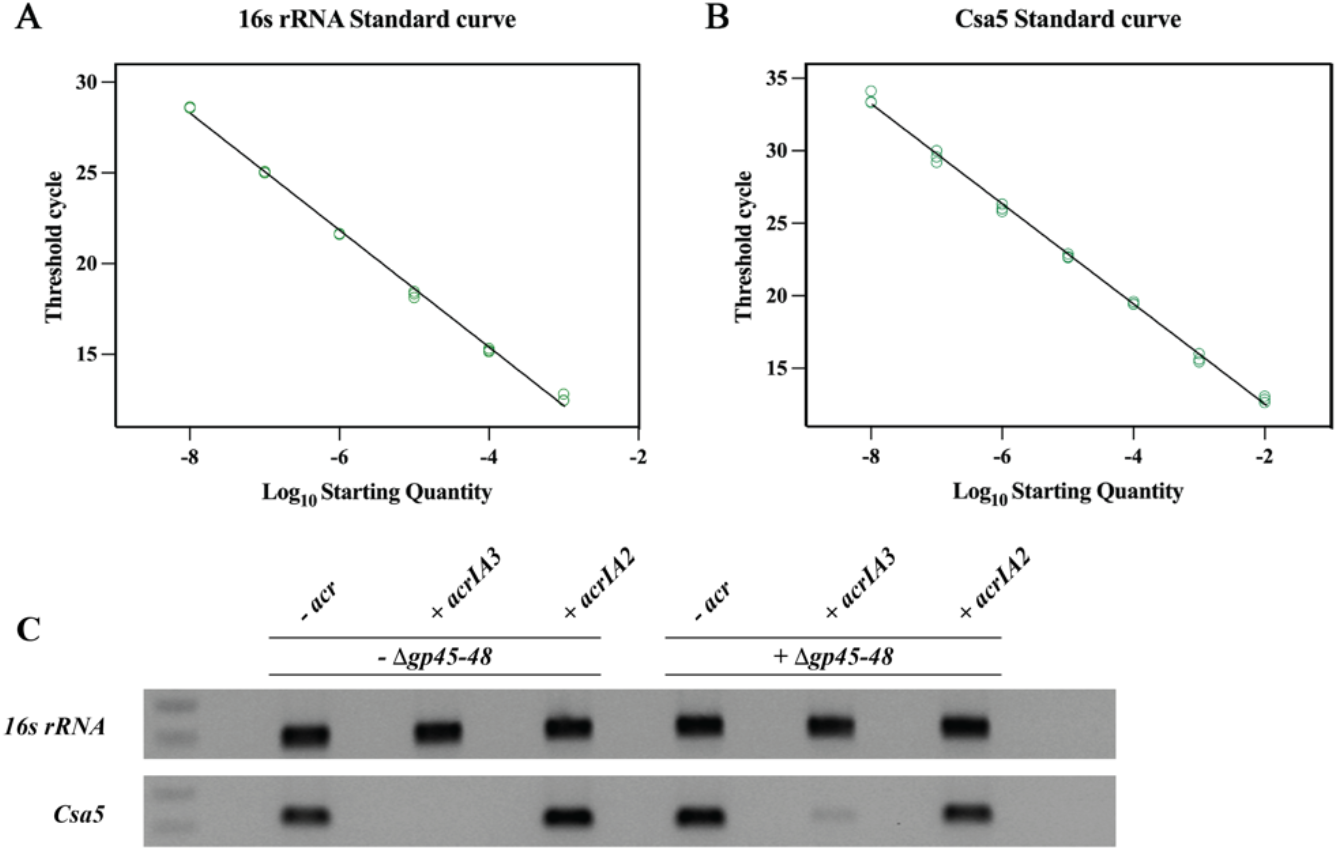
Determination of qPCR efficiencies for primer sets matching *16sRNA* (A) and *csa5* (B). The standard curve was constructed using templates based on *16sRNA* fragment and *csa5* gene cloned into *E. coli* plasmids. The qPCR efficiencies for the standard curves of *16srRNA* and *csa5* were 104.17% and 94.80% respectively. **C**. qRT-PCR products from Figure 2 were separated by gel electrophoresis on a 1.2% Agarose gel.

**Supplementary figure 3.**
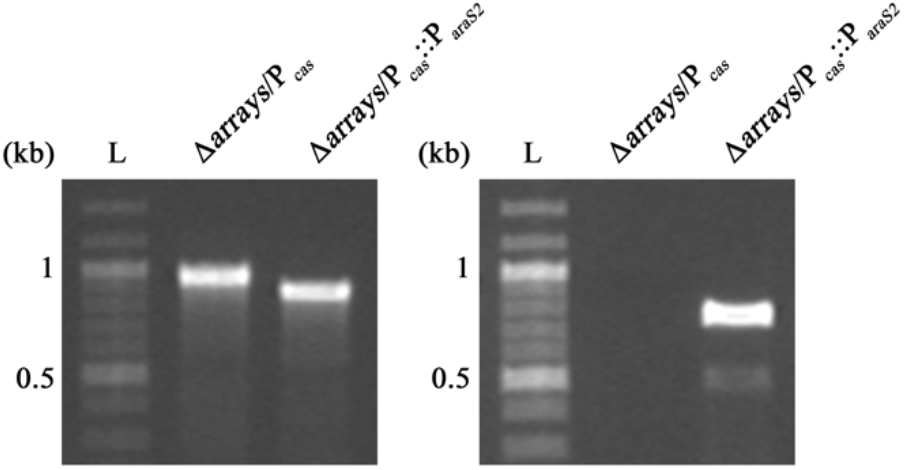
Verification of Δ*arrays*/P_*cas*_::P_*araS2*_ strain. PCR analysis of Δ*arrays*/P_*cas*_::P_*araS2*_ and its parental strain Δ*arrays*/P_*cas*_ with primers flanking the promoter (P_*cas*_) region (Left panel). Another primer annealing specifically to the arabinose promoter was used in combination with a primer specific for subtype I-A system encoded on *S. islandicus* (Right panel). L, log-2 DNA ladder.

**Supplementary figure 4.**
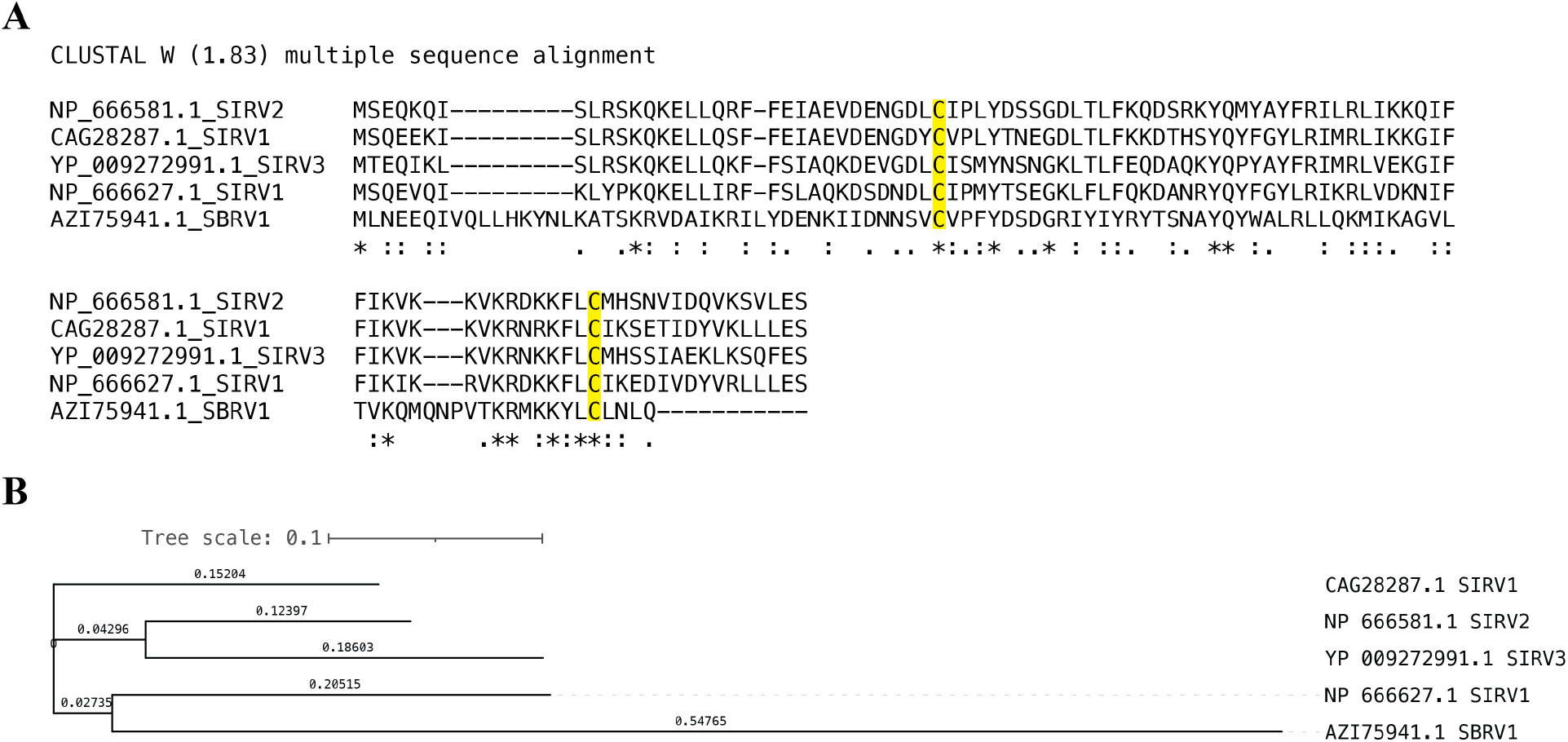
Multiple sequence alignment and phylogenetic analysis of SIRV2 gp47/AcrIA3. **A**. Multiple sequence alignment of SIRV2 gp47 (NP_666581.1) homologs as identified by PSI-BLAST analysis. **B**. Phylogenetic tree of SIRV2 gp47 homologs. GeneBank identifiers of individual sequences are indicated along with viral strains on which they are encoded.

**Supplementary figure 5.**
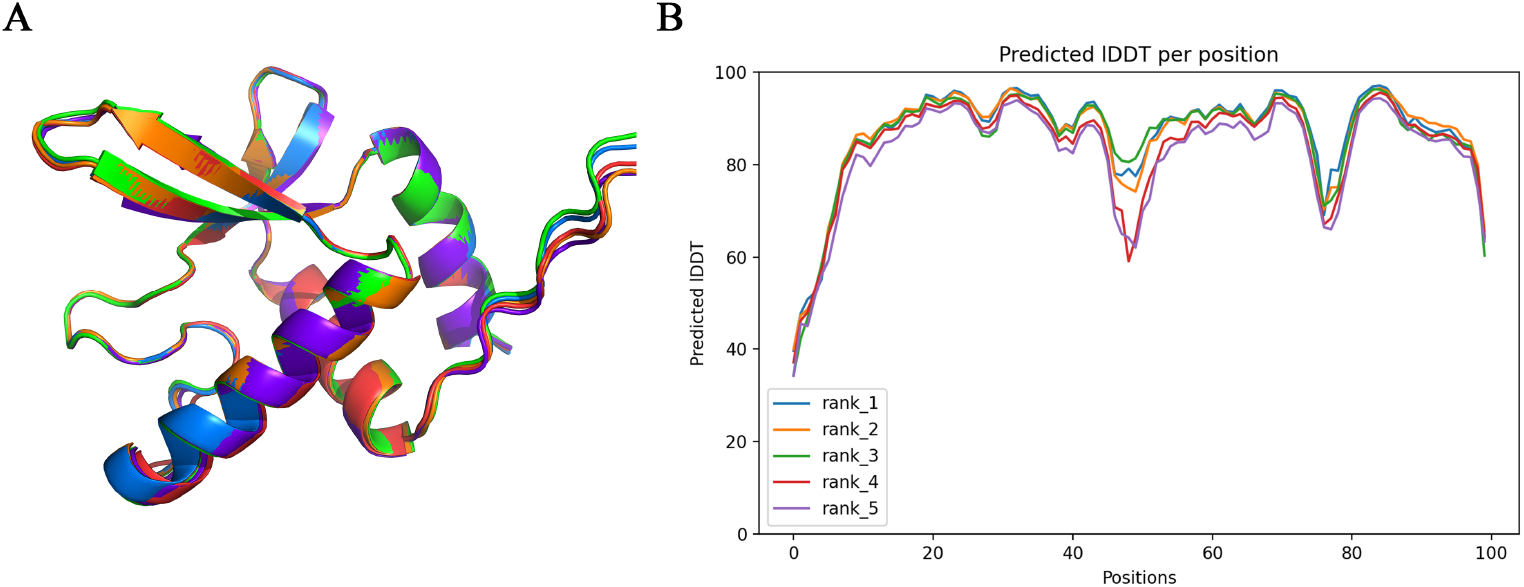
AlphaFold2 model prediction. **A**. Superimposition of the 5 models predicted by AlphaFold2. **B**. pLDDT, per-residue estimate of confidence in the structures predicted by AlphaFold2.

**Supplementary table 1:**
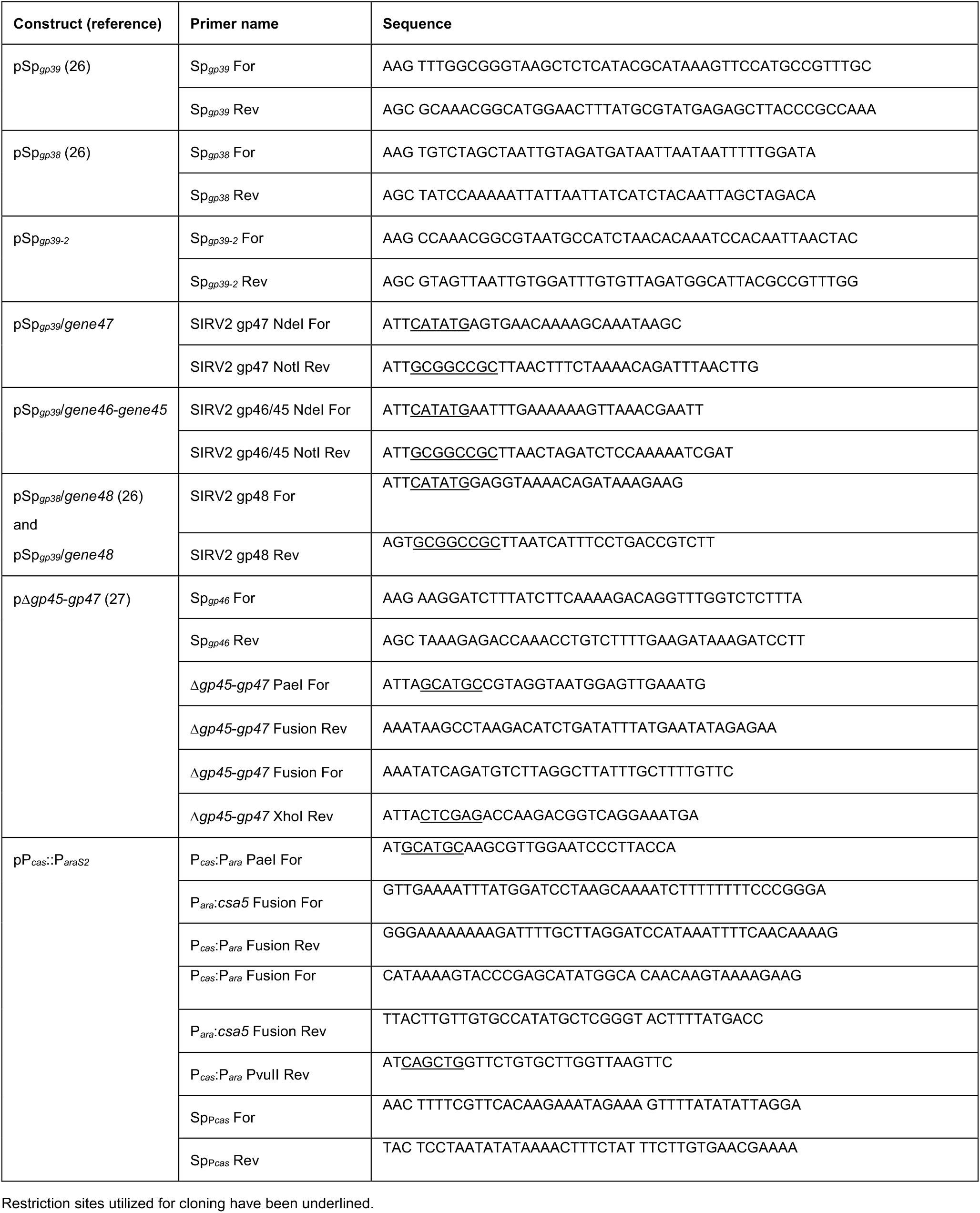
List of primers used in this study.

